# HU promotes higher-order chromosome organisation and influences DNA replication rates in *Streptococcus pneumoniae*

**DOI:** 10.1101/2024.09.27.615122

**Authors:** Maria-Vittoria Mazzuoli, Renske van Raaphorst, Louise Martin, Florian Bock, Agnès Thierry, Martial Marbouty, Barbora Waclawikova, Jasper Stinenbosch, Romain Koszul, Jan-Willem Veening

## Abstract

Nucleoid-associated proteins (NAPs) are crucial for maintaining chromosomal compaction and architecture and are actively involved in DNA replication, recombination, repair, and gene regulation. In the opportunistic pathogen *Streptococcus pneumoniae,* HU is the only identified NAP, and its role in chromosome conformation and other essential processes has not yet been investigated. Here, we use a multi-scale approach to explore the role of HU in chromosome conformation and segregation dynamics. By combining superresolution microscopy and whole-genome binding analysis, we describe the nucleoid as a dynamic structure where HU binds transiently across the entire nucleoid, with a preference for the origin of replication over the terminus. Reducing cellular HU levels impacts nucleoid maintenance and disrupts robust nucleoid scaling with cell size. This effect is similar to the distortion caused by fluoroquinolone-antibiotics, supporting earlier observations that HU is essential for maintaining DNA supercoiling. Furthermore, in cells lacking HU, the replication machinery is misplaced, and cells are unable to initiate and proceed with on-going replication. Chromosome conformation capture (Hi-C) experiments revealed that HU is required to maintain cohesion between the two chromosomal arms, in a similar way to the structural maintenance of the chromosome complex SMC. Together, we show that by promoting long-range chromosome interactions and supporting the architecture of the domain encompassing the origin, HU is fundamental for chromosome integrity and the intimately related processes of chromosome replication and segregation.

**Graphical abstract:** 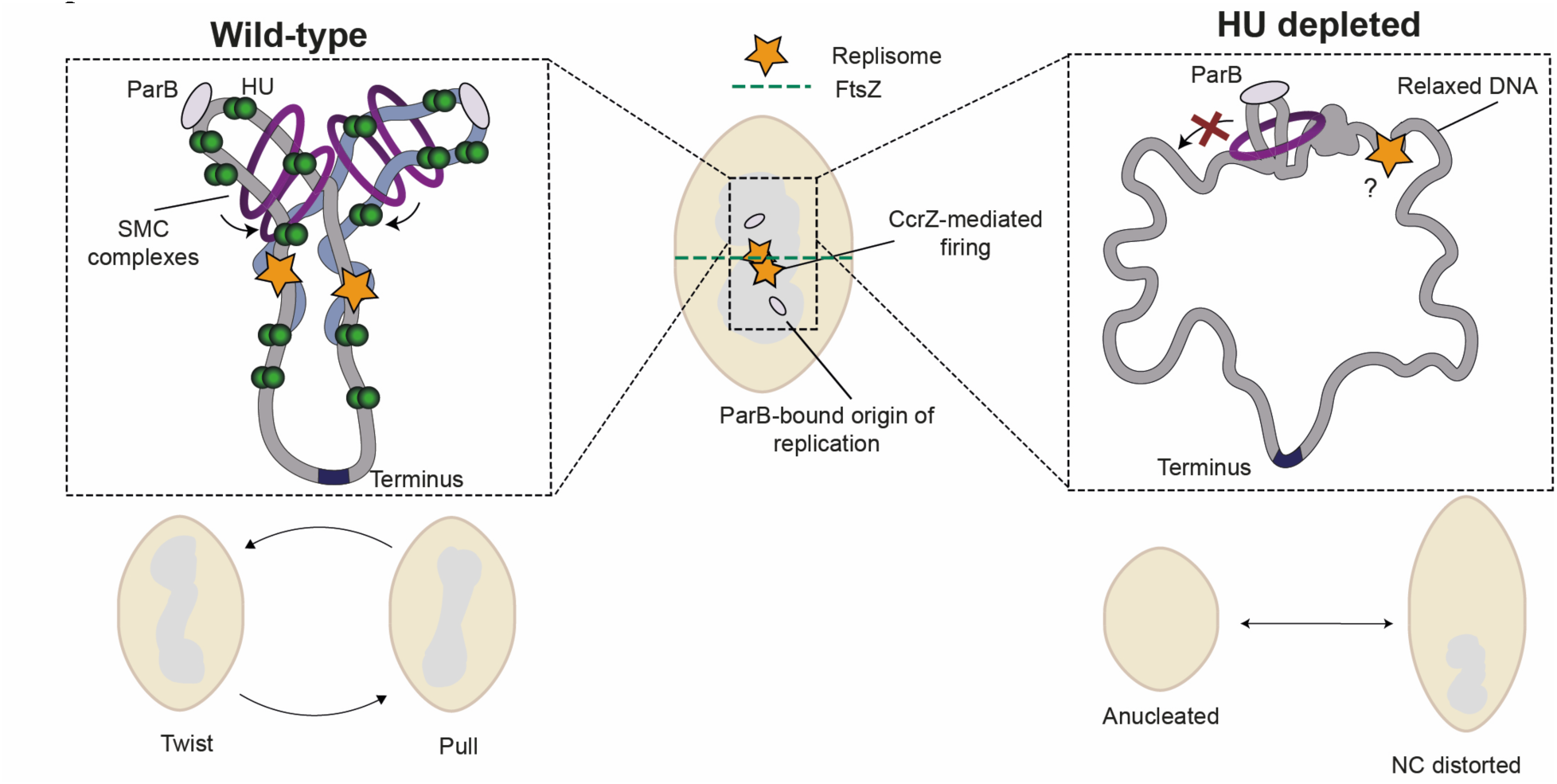

## Introduction

In bacteria, unlike eukaryotes, DNA replication, segregation and compaction occur simultaneously. Rather than a lack of organisation, this points to an intimate connection and tight control of all these processes during the bacterial cell cycle. Chromosome compaction is mediated by several factors, including DNA supercoiling (Dame et al. 2020), transcription (Bignaud et al. 2024), protein complexes of the structural maintenance of chromosome (SMC) family (Mäkelä and Sherratt 2020), and nucleoid-associated proteins (NAPs) (Browning, Grainger, and Busby 2010). The degree of supercoiling, the abundance of NAPs and condensin complexes, along with their impact on chromosome structures, largely differ among bacteria (Ponndara et al. 2024). Notably, the pneumococcus possesses only the α subunit of HU (α-HU), which is highly expressed and crucial for growth (Ferrándiz et al. 2018, X. Liu et al. 2017), whereas most bacterial species have multiple nucleoid-associated proteins (NAPs) from different classes. Chromosome conformation capture coupled to deep sequencing (Hi-C) has provided detailed topological insights into the architecture of the genome within the cell of well-studied bacterial models (Badrinarayanan, Le, and Laub 2015; Hofmann and Heermann 2018; Yáñez-Cuna and Koszul 2023). Interestingly, the effects of HU loss on cell survival and large-scale structuring seems to differ widely between species (Ponndara et al. 2024). Removing HU in *E. coli* causes reduced viability, while it is lethal in *Bacillus subtilis* (Micka et al. 1991; Huisman et al. 1989). Hi-C showed that *E. coli Δhupα/β* mutants are deficient in long-range DNA interactions and show an increase in short-range interactions (Lioy et al. 2018). Superresolution imaging of enlarged, circular *E. coli* confirms that large subdomains are substituted for smaller subdomains in *Δhupα/β* mutants (Wu et al. 2019). On the opposite, in *Caulobacter crescentus*, removing HU promotes short-range contacts (Le et al. 2013). Other than its structural role, HU is directly involved in essential cellular processes, including replication (Chodavarapu et al. 2008), recombination and repair (Kamashev and Rouviere-Yaniv 2000), and gene regulation (Oberto et al. 2009; Stojkova et al. 2018; Mangan et al. 2011). Recently, HU was shown to be required for replication initiation in *B. subtilis* and *Staphylococcus aureus* (Karaboja and Wang 2022; Sharkey et al. 2023; Schramm Frederic D. and Murray Heath 2022) and for the organisation of the origin of replication in *Mycobacterium tuberculosis* (Hołówka et al. 2017). Pneumococcal HU was shown to be required to maintain DNA supercoiling (Ferrándiz et al. 2018), together with the DNA topoisomerase I (TopA), (de la Campa et al. 2017) and its regulator StaR (de Vasconcelos Junior et al. 2023).

Over the past decade, *S. pneumoniae* has become a key model organism for uncovering the molecular mechanisms that regulate the bacterial cell cycle. However, the involvement of HU in supporting different phases of the cell cycle has not yet been studied. In the pneumococcus, DNA replication occurs at mid-cell (Slager et al. 2014) and is orchestrated at the replication origin (*oriC*) by DnaA, whose activity is regulated by CcrZ (Gallay et al. 2021). Additionally, YabA negatively regulates initiation of DNA replication (Felicori et al. 2016, Gallay et al. 2021). Besides the active process of DNA replication, another mechanism important for driving chromosome segregation is the *parS*/ParB system. ParB (partial partition system B) binds to *parS* sites near the replication origin, recruiting SMC to ensure proper chromosome segregation (Minnen et al. 2011). RocS, a membrane-binding protein, interacts with ParB and localizes at the origin, and is required for segregation (Mercy et al. 2019). FtsK, which functions as a motor to pull through nearly segregated DNA in *E. coli* (Stouf, Meile, and Cornet 2013), might also be required for chromosome segregation in *S. pneumoniae* (X. Liu et al. 2017; Le Bourgeois et al. 2007). Supercoiling and transcription also modulate chromosome segregation (Kjos and Veening 2014). Surprisingly, most of the factors required for this process, such as ParB and SMC, can be deleted with only mild phenotypes (van Raaphorst, Kjos, and Veening 2017; Kjos and Veening 2014), suggesting that other factors or processes required for active segregation are yet to be elucidated.

Here, we employ a multi-scale approach of live-cell imaging and genome-scale analysis to dissect the role of HU in chromosome conformation, replication, and segregation. Tracking the pneumococcal nucleoid with HU using superresolution microscopy showed that it is extremely compact and dynamic. While HU binds transiently the whole chromosome, ChIP-Seq analysis revealed a binding preference in proximity of the origin of replication. By depleting HU, we observed a drastic loss of nucleoids, aberrations in cell size, and, consequently, a loss of scale between nucleoids and cell size. Quantification of DNA replication rates and integration of Hi-C data with fluorescence and superresolution microscopy provided a model in which HU-mediated structural (both short- and large-scale) and supercoiling defects are deleterious to the progression of the replication machinery and successful chromosome segregation.

## Materials and methods

### Bacterial strains and growth conditions

All strains, plasmids and primers used are listed in Supplementary Table 1 and Supplementary Table 2. All pneumococcal strains in this study are derivatives of *S. pneumoniae* D39V and are listed in Supplementary Table 1. Construction of strains is described in the Supplementary information. Strains were grown in liquid C+Y medium at 37 °C from a starting optical density (OD600) of 0.01 until the appropriate OD. Induction of the anhydrotetracycline-inducible promoter (*Ptet*) was carried out by supplementing the medium with 25 ng/ml aTc (Sigma-Aldrich) and the IPTG-inducible promoter (*Plac*) with 1 mM IPTG (Sigma-Aldrich). Depletion strains were first grown in presence of 25 ng/ml aTc up to OD600 = 0.1, then washed three times in fresh medium, diluted 15 times in fresh medium and grown until the desired OD. Transformation of *S. pneumoniae* was performed as described previously (Domenech, Slager, and Veening 2018) with cells taken at the exponential growth phase (OD600 = 0.1). When necessary, the medium was supplemented with the following antibiotics: chloramphenicol (4 µg ml−1), erythromycin (0.5 µg ml−1), kanamycin (250 µg ml−1), spectinomycin (100 µg ml−1) and tetracycline (0.5 µg ml−1).

### SIM microscopy

Samples were prepared as for time-lapse wide field microscopy, with the exception that for snap shots, the C+Y medium was substituted for PBS, and a high precision 17µm coverslip was used (VWR). 3D SIM experiments were performed on a Deltavision OMX SR (GE Healthcare) microscope equipped with 488 nm and 568 nm lasers. 9-12 Z-stacks of 125 nm distance were acquired in 3D-SIM mode. For time lapse imaging, a 3D-stack was taken every 10 minutes, while a constant temperature of 37°C was maintained. Reconstruction and channel alignment was done in SoftworX (GE Healthcare) with ‘force modulation amplitude’ checked on and furthermore standard settings.

### PALM microscopy and analysis

PALM microscopy was performed on a Deltavision Elite microscope equipped with DLM (Deltavision Localization Microscopy), using a 60x oil 1.49 NA TIRF objective (Olympus), 405 nm, 588 and 568 nm lasers, a sCMOS camera. PALM imaging was done in Hi-Lo mode at 37°C. Cells were prepared as for Wide-Field time-lapse microscopy, where C+Y medium was substituted with PBS. Before each acquisition, a green fluorescence image (excitation: 488 nm laser, emission 525/50 nm) and DIC image of the cells was made. After this, PALM images were acquired in the following sequence: activation (405 nm) – 4x red excitation (568 nm), while red emission (632/60 nm) was recorded. The activation power was set to very low in the beginning of the acquisition and manually increased over time. 5000 images were taken for each PALM acquisition.

### Particle tracking

To determine the mean square displacement (MSD) of HU-meos3.2, iSBatch (Caldas et al., 2015) was used to track HU-meos3.2 in the images acquired using PALM. For this, firstly the regions containing the cells were selected in imageJ, after which the Peak Fitter was used to find the fluorescent spots. Using the Particle Tracker, allowing for distances travelled not longer than 4 pixels/frame, only the spots appearing in multiple frames were kept. Tracks longer than 15 frames were removed. Finally, the collective MSD was calculated. This procedure was repeated for 10 PALM movies. All MSD tables were collected and plotted in R.

### Phase contrast and fluorescent microscopy

*S. pneumoniae* cells were grown in C+Y medium pH 7.4 at 37 °C to an OD600 = 0.1 with 25 ng/ml aTc at 37 °C, after which they were washed three times with fresh C+Y medium, with or without the inducers when appropriate: IPTG for activation of dCas9 or complementation with fluorescent fusions (mCherry) or aTc for HU expression in depletion strain. 1 ml of culture at OD600 = 0.1 was harvested by centrifugation for 1 min at 9,000*g*. For DAPI staining, 1 µg ml−1 of DAPI (Sigma-Aldrich) was added to the cells and incubated for 2 min at room temperature before centrifugation. For imaging of exponentially growing cultures, cells were washed twice with 1 ml of ice-cold PBS and resuspended in 50 µl of ice-cold PBS. One microliter of cells was then spotted onto PBS-pads. Pads were then placed inside a gene frame (Thermo Fisher Scientific) and sealed with a cover glass as described previously (de Jong et al. 2011). Microscopy acquisition was performed using either a Leica DMi8 microscope with a sCMOS DFC9000 (Leica) camera and a SOLA light engine (Lumencor) or a DV Elite microscope (GE Healthcare) with a sCMOS (PCO-edge) camera and a DV Trulight solid-state illumination module (GE Healthcare), and a ×100/1.40 oil-immersion objective. Phase-contrast images were acquired using transmission light (100 ms exposure). Fluorescence images were usually acquired with 700 ms exposure. The Leica DMi8 filters set used were as followed: DAPI (Leica 11533333, Ex: 395/25 nm, BS: LP 425 nm, Em: BP 460/50 nm), GFP (Ex: 470/40 nm Chroma ET470/40x, BS: LP 498 Leica 11536022, Em: 520/40 nm Chroma ET520/40 m), and mCherry (Chroma 49017, Ex: 560/40 nm, BS: LP 590 nm, Em: LP 590 nm). Images were processed using LasX v.3.4.2.18368 (Leica).

### Cell and nucleoid segmentation and analysis

Cells and nucleoids imaged using SIM were segmented using Morphometrics (Ursell et al. 2017). Cells and nucleoids imaged using phase-contrast/wide-field fluorescence microscopy was segmented using Oufti (Paintdakhi et al. 2016), where the nucleoids were segmented in object detection mode. DnaX foci where detected using the Peak Fitter tool from ISBatch (Caldas et al. 2015) To determine the cell and nucleoid dimensions in the PALM experiments, cell outlines were drawn by hand three times from DIC images (cells) and PALM reconstructions (nucleoids) using FIJI (Schindelin et al. 2012) and subsequently averaged. After segmentation, the wide-field and SIM microscopy data was analysed and visualised in R (https://cran.r-project.org/) using the R packages BactMAP (van Raaphorst, Kjos, and Veening 2020) and ggplot2 (Wickham 2016). To determine nucleoid size, dimensions and intensity, firstly, the intensities from the TIFF values were added to the segmented nucleoids using BactMAP::extr_OriginalCells(). This returned the dimensions and size of the nucleoids and a list of pixel values per nucleoid. Of these values, the median and standard deviation was determined. The cell dimensions were imported from the Oufti or Morphometrics segmentations.

### Microtiter plate-based growth assay

For *S. pneumoniae* growth assays, cells were first grown in C+Y medium until the mid-exponential growth phase (OD600 = 0.3) with the inducer at 37 °C, after which they were washed three times with fresh C+Y medium, with or without the inducers (IPTG or aTc) when appropriate. Cellular growth was then monitored every 10 min at 37 °C in a microtiter plate reader (TECAN Infinite F200 Pro). Each growth assay was performed at least in triplicate. The average of the triplicate values was plotted, with the s.e.m. represented by an area around the curve.

### *ori/ter* ratio determined by whole-genome sequencing

Cells were pre-grown until OD600 = 0.3 in C+Y medium at 37 °C in presence of the inducer (aTc) for HU depletion strain (VL5898), and in CY only for WT D39V. Cells were then washed three times with CY and diluted 15 times in 10 ml of fresh C+Y medium supplemented with or without the inducer and harvested for genomic DNA isolation after one, two and three hours of depletion (OD600 = 0.1-0.2). Genomic DNA was isolated using the FastPure Bacteria, DNA isolation Mini Kit (Vazyme). Samples were sequenced with Illumina PE150 by Novogene, and, on average, 14,530,047 raw reads (maximum: 24,200,092 - minimum 12,279,628). Raw Illumina sequencing reads were checked for quality using FastQC v0.11.8, available from https://www.bioinformatics.babraham.ac.uk/projects/fastqc/, and aligned to the D39V genome using Bowtie 2.0 (Langmead and Salzberg 2012). Alignment files were used as input for iRep (available at https://github.com/christophertbrown/iRep) (Brown et al. 2016).

### Western blot analysis

Cells were grown in C+Y medium until OD600 = 0.1 and harvested by centrifugation at 8,000*g* for 2 min at room temperature from 1 ml of culture. Cells were resuspended in 150 µl of Nuclei Lysis Solution (Promega) containing 0.05% SDS, 0.025% deoxycholate and 1% protease inhibitor cocktail (Sigma-Aldrich) and incubated at 37 °C for 20 min and at 80 °C for 5 min to lyse the cells. One volume of 4× SDS sample buffer (50 mM Tris–HCl pH 6.8, 2% SDS, 10% glycerol, 1% β-mercaptoethanol, 12.5 mM EDTA and 0.02% Bromophenol blue) was then added to three volumes of cell lysate sample and heated at 95 °C for 10 min. Protein samples were separated by SDS–PAGE (4–20%) and blotted onto polyvinylidene fluoride membranes (Merck Millipore). Membranes were blocked for 1 h with Tris-buffered saline (TBS) containing 0.1% Tween 20 (Sigma-Aldrich) and 5% dry milk and further incubated for 1 h with primary antibodies diluted in TBS, 0.1% Tween 20, 5% dry milk. Commercial polyclonal rabbit anti-GFP IgG (Invitrogen A-6455) was used at 1:5,000. Membranes were washed four times for 5 min in TBS, 0.1% Tween 20 and incubated for 1 h with secondary goat anti-rabbit IgG horseradish peroxidase conjugated (Abcam AB205718) diluted 1:20,000 in TBS, 0.1% Tween 20 and 5% dry milk. Membranes were then washed four times for 5 min in TBS, 0.1% Tween 20 and revealed with Immobilon Western HRP Substrate (Merck Millipore).

### ChIP-Sequencing and analysis

Cells were grown in 2 mL of C+Y, 37°C up to OD_600_ ∼ 0.4, diluted 50 times in 30 mL of preheated C+Y (37°C) and allowed to grow up to an OD_600_ of 0.2. The cells were harvested and fixated in 1% formaldehyde. The cells were washed several times in ice-cold PBS and the pellets were snap-frozen and stored in -80°C. Subsequently, the cells were thawed and sheared in a Covaris S2 sonicator using the following protocol: 200 cycles, 100 W, 10% load, 12 min. A fraction of the cell mixture was taken apart (input fraction). Polyclonal anti-GFP (ThermoFisher A-6455) was bound to magnetic beads (Dynabeads Protein G 10004D). The remaining sheared cells (IP fraction) were added to the beads and the mixture was incubated for two hours on a rotating wheel for two hours. After immunoprecipitation, the beads were washed. Cross-linking was reversed in both the input and IP fraction by shaking 1400 rpm overnight at 65°C in 1% SCS, 10 mM EDTA and 50 mM Tris (pH 8). DNA was purified using phenol-chloroform extraction followed by a PCR purification kit (Qiagen QIAquick). Libraries were prepared by the Lausanne Genomic Technologies Facility (LGTF) using an Ovation Ultralow V2-DNA-Seq library preparation kit (NuGEN) and 100 nt paired-end sequencing was done at the LGTF on an Illumina HiSeq. For the analysis, sequence trimming, alignment and peak annotation was done using a snakemake pipeline (https://github.com/veeninglab/chip-seq). Sequence files were gunzipped and sequence quality was assessed using FastQC. Data was trimmed using Trimmomatic (Bolger, Lohse, and Usadel 2014) and sequences were aligned to the reference genome (D39V, GenBank CP027540) using Bowtie 2.0 (Langmead and Salzberg 2012) and converted to *.bam* files using SamTools (Li et al. 2009) and filtered to remove false positive peaks due to duplicated regions in the chromosome using Sambamba (Tarasov et al. 2015). Peaks were annotated using MACS2 (T. Liu 2014). Sequence read depths were binned by 500 bp and normalised per sample. Relative enrichment was plotted in R by dividing the normalised input over the normalised IP. The relative enrichment of HU per binned region (1000 bps) was plotted against RpoB enrichment and GC content (0-1) per 1000 base pairs.

### Hi-C procedure and sequencing

Cells were pre-grown until OD600 = 0.3 in C+Y medium at 37 °C in presence of the inducer (aTc) for HU-depletion strain (VL5898). Cells were then washed three times with CY and diluted 15 times in 10 ml of fresh C+Y medium supplemented with or without the inducer and let grow for three hours up to OD600 = 0.1 - 0.2. Cell fixation was performed with 3% formaldehyde (Sigma-Aldrich, cat. no. F8775). Quenching of formaldehyde with 300 mM glycine was performed at room temperature for 20 min under gentle agitation. The samples were recovered by centrifugation (4000 g, 10 min, 4 °C), washed with 10 mL 1X PBS, re-centrifuged, and stored at − 80 °C until processing. Hi-C experiments were performed as described in ref. (Cockram, Thierry, and Koszul 2021), as well as DNA extraction, purification, and processing into a sequencing library. Proximity ligation libraries were sequenced using pair-end (PE) Illumina sequencing (2 × 35 bp, NextSeq500 apparatus) at the sequencing platform of the Institut Pasteur of Paris.

### Processing of reads and Hi-C data analysis

Reads were aligned to the reference genome with Bowtie 2.0 and Hi-C contact maps were generated using hicstuff v3.1.5 (Matthey-Doret et al. 2022), employing default settings and DpnII and HinfI enzymes for digestion. Contacts were filtered according to the method described in ref. (Cournac et al. 2012), and PCR duplicates, identified as paired reads mapping to the exact same position, were removed. The resulting matrices were binned at intervals of 1, 5, 10, and 20 kb. Balanced normalisation was achieved using the ICE algorithm (Imakaev et al. 2012) and the contact maps were stored in cool file format via cooler (v0.8.11) (Abdennur and Mirny 2020). Comparative analyses were conducted by binning matrices at 5 kb resolution, downsampling to equal contact numbers, and comparing using the log2 ratio.

## Results

### The pneumococcal nucleoid and HU are highly dynamic

Because of its very high abundance and limited cellular diffusion, fluorescent protein fusions to HU are widely used to track live pneumococci in animal and tissue models as well as flow cytometry studies (Kjos et al. 2015; Dewachter et al. 2022; Pulous et al. 2022; Ercoli et al. 2018). In other bacteria, HU has widely been used as a tool for live-cell tracking and measurement of nucleoid dimensions and dynamics (Bettridge et al. 2021; Floc’h et al. 2019; Fisher et al. 2013). To test whether *S. pneumoniae* HU is an appropriate proxy for the pneumococcal nucleoid, we performed co-localization studies using HU-FP fusions and DAPI. As shown in Fig. 1A, we observed near-perfect overlaps demonstrating that HU is a suitable marker for the pneumococcal nucleoid. This enables us to investigate the shape and dynamics of the nucleoid during the cell cycle in live *S. pneumoniae* cells.

**Figure 1.**
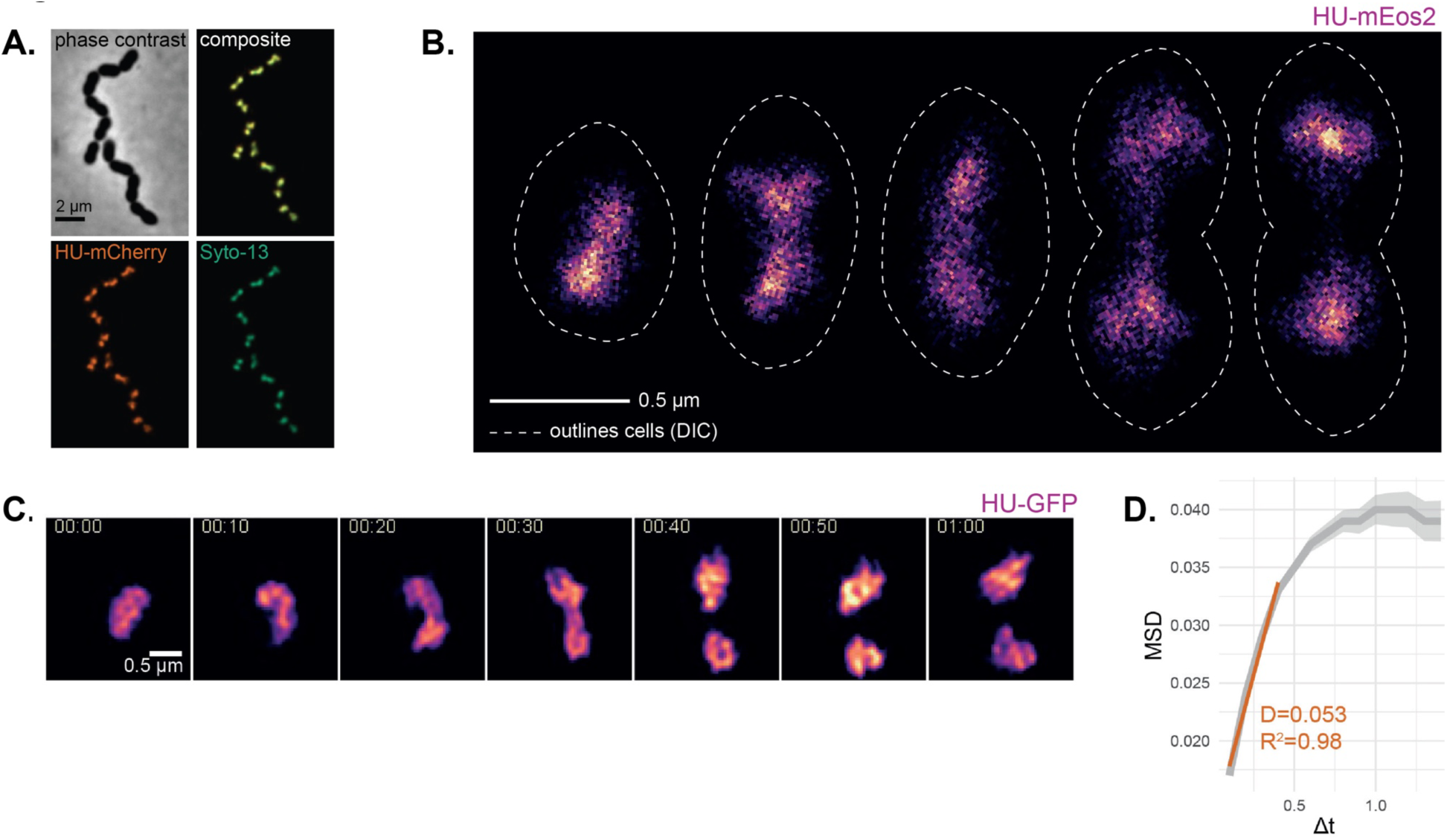
A dynamic, compact nucleoid robustly scales with cell size. **A.** Deconvolved epi-fluorescence microscopy images of HU-sfGFP and the chromosome stained with DAPI show a near-perfect overlap. **B.** Representative images of PALM microscopy reconstructions of 5 cells with increasing cell sizes. PALM reconstructions of experiment detecting HU-mEos3.2. Scale bar: 0.5 μm. Dashed lines: cell outline, drawn by hand from DIC images. **C.** Time-lapse sequence of HU-sfGFP imaged using structured illumination microscopy, mid-section of 3D-SIM experiment. Image taken every 10 minutes, scale bar 0.5 micron. **D.** Mean Square Displacement (MDS) over time of HU-meos3.2. HU-meos3.2 recorded using PALM, particles were tracked, and collective MSD was calculated using iSBatch (Caldas et al. 2015). Apparent diffusion coefficient for the short time intervals was calculated by determining the linear slope between 0-0.5 s. Shade=standard error.

To analyse the pneumococcal nucleoid in high temporal and spatial resolution in live cells, we performed time-lapse structured illumination microscopy (SIM) of cells carrying a functional fusion of HU to superfolder GFP (HU-sfGFP) (Supplementary Fig. 1A). This showed that pneumococcal nucleoids are very dynamic, changing shape and structure over time (Fig. 1B). The bulk nucleoid splits at the end of the cell cycle (40 min), which is confirmed by the observation that 96% of single pneumococcal cells carry one bulk nucleoid (Supplementary Fig. 1B and C).

While SIM provides higher spatial resolution compared to wide field epifluorescence microscopy, since the pneumococcal nucleoid has a diameter of ∼0.5 µm, we are imaging close to the resolution limit. To obtain deeper insights into the pneumococcal nucleoid shape in live cells, we set up Photoactivated Localization Microscopy (PALM) for *S. pneumoniae*. To perform PALM, we first constructed *S. pneumoniae* carrying HU fused to a codon-optimised photoswitchable mEOS3.2 (Zhang et al. 2012). In total, 160 pneumococcal cells carrying hu-meos3.2 were imaged, with a Nyquist resolution of 18 nm and an average number of 3.990 ± 2.730 HU dimers per cell. Cells were divided in five groups based on cell length. Consistently with the observations of the SIM microscopy, bulk nucleoids stretch out to the new mid-cell regions early in the cell cycle, but only the largest cells carry two separated nucleoids. Interestingly, in the second-largest cell group, the region between the dense domains of the nucleoid became highly stretched measuring only 163 ± 45 nm, which is beyond the resolution limits of our SIM microscope. Not only is the nucleoid very dynamic as shown by SIM (Fig. 1C and 1D), PALM shows that HU itself is also dynamically moving within the nucleoid space with a D_app_ of 5.3*10-2 µm^2^s^-1^ measured over short time intervals (< 0.5 s). This is more than 20-fold more mobile than ParB_L.lactis_-GFP bound to a single *parS_lactis_*-site at the origin of replication, measured previously using TIRF microscopy (van Raaphorst et al., 2017), and 10-fold more mobile than DnaX-GFP, however, more confined than free diffusing GFP (Eldar and Elowitz 2010), indicating that HU is transiently binding the nucleoid.

### HU display low intensity binding with bias toward the origin and highly transcribed loci

To assess the properties of HU binding at a whole-genome scale, we performed Chromatin ImmunoPrecipitation-sequencing (ChIP-seq) of pneumococcal cells carrying HU-sfGFP. Two independent cell cultures of exponentially growing bacteria were used for immunoprecipitation with an anti-GFP antibody, and non-immunoprecipitated samples were used as controls. After sequencing and bioinformatic analysis, we detected 367 reproducible high confidence binding sites (qvalue < 0.05) of enrichment along the entire chromosome, which are absent in non-immunoprecipitation controls (Fig. 2A). These peaks are wide-spread and have a relatively weak signal (log 2 fold changes between 1 and 2), similarly to what has been observed for *E. coli* α/b HU subunits (Prieto et al. 2012). Interestingly, we detected only a few binding sites (35 out of 367) near the replication terminus (*ter*) (1-1.04 Mbp), indicating a bias in HU binding distribution along the chromosome, with a preference for the origin *(ori)* region (Supplementary Table 3). Since ChIP-Seq sequencing data was adjusted using input DNA from cells in the same growth stage, we can exclude that this effect is due to the impact of replication-associated gene dosage effects typically seen during exponential growth. Additionally, peak lengths are variable and can reach up to 7 kb, indicative of a dispersed HU binding. Despite the relatively low enrichment levels measured that may be caused because of its dynamic ON/OFF binding (Fig. 1D), the high reproducibility of the distribution patterns and their detection with a strict statistical threshold (using q-value instead of p-value) resulted in binding profiles that are representative of HU.

**Figure 2.**
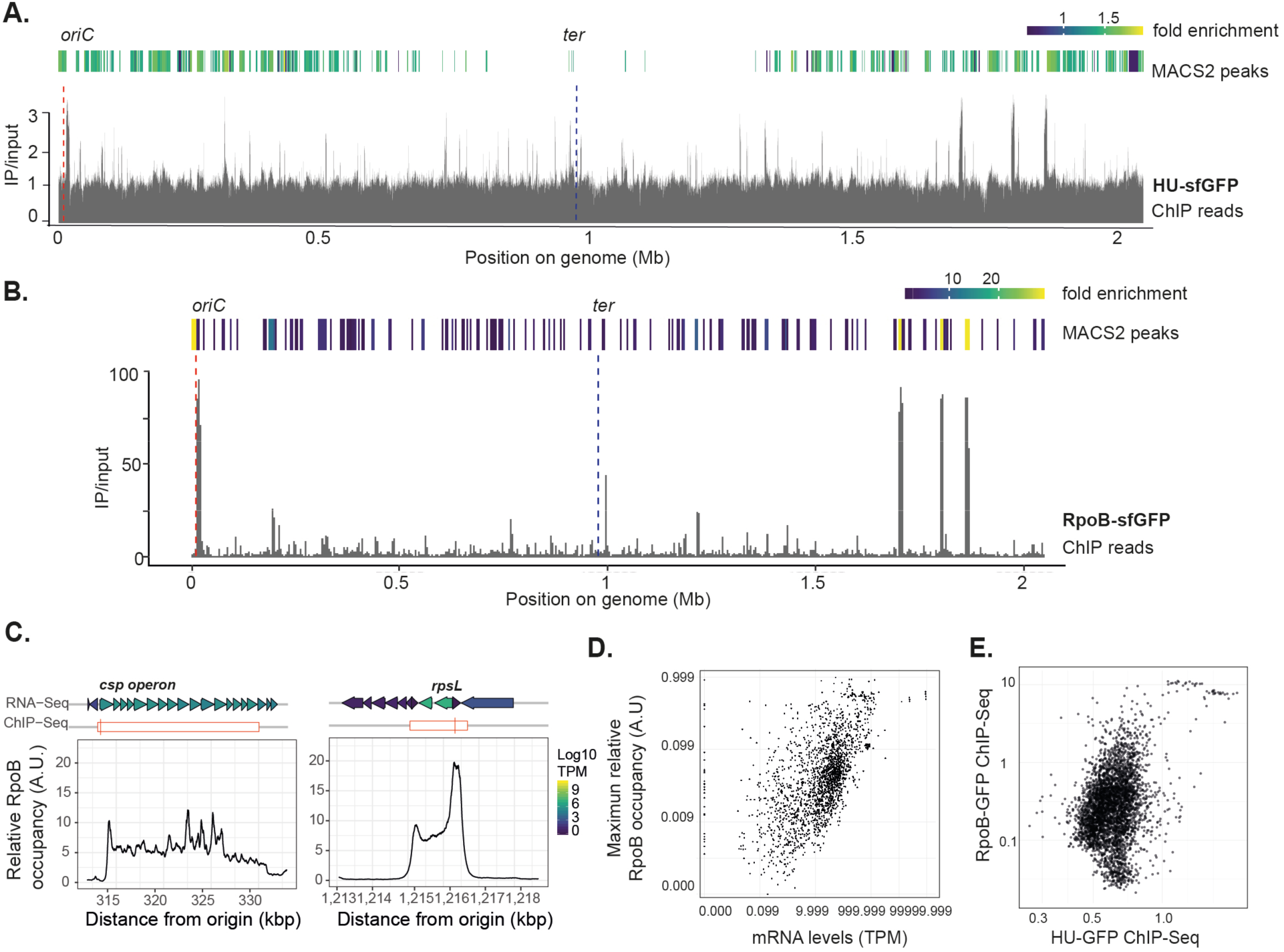
HU binding profile and correlation with active transcription. **A.** HU binding profile obtained by ChIP-Seq. Normalised counts (IP/input) binned per 50 base pairs obtained for the immunoprecipitated HU-sfGFP sample over the input (non-immunoprecipitated) sample are plotted against the chromosomal coordinates. Significant (qvalue < 0.05) peaks detected by MACS2 are depicted, and peaks with a fold change > 2 are coloured yellow. Position of origin and terminus of replication are indicated on top of the graph. Note the asymmetric distribution of the left and right arms of the chromosome due to a chromosomal inversion in strain D39 (Slager et al 2018). **B.** Enrichment profile of ChIP-Seq for RpoB-sfGFP (normalised IP/input per 50 base pairs). **C**. Relative RpoB occupancy at highly transcribed regions (*csp*, capsule operon and *rplS*, ribosomal RNA). Absolute MACS2-estimated summit indicated with vertical red bar. Gene expression levels are depicted by colours. **D.** RpoB-GFP occupancy sites per gene vs mRNA levels in C+Y (Aprianto et al. 2018)**. E.** RpoB vs HU occupancy (IP/input reads normalised with total reads).

Transcription imposes structural constraints on chromosome folding (Bignaud et al. 2024) and could therefore affect HU binding pattern. To test whether there is correlation between HU binding sites and active transcription, we performed ChIP-seq on cells carrying a functional GFP-marked β-subunit of RNA polymerase (RpoB-sfGFP) (Fig. 2B and Supplementary Table 3). Magnification of highly transcribed genes such as the capsule operon *cps* and the ribosomal gene *rpsL* showed a high median RpoB occupancy at these chromosomal locations (Fig. 2C). As expected, comparison of RpoB occupancy and gene expression, quantified as the average of Transcripts Per Million (TPM) per gene (Aprianto et al. 2018), showed positive correlation (Fig. 2D). After removal of ribosomal RNA and tRNA loci, we highlighted a slight correlation between HU and RpoB binding sites, suggesting that more HU binding occurs in proximity of actively transcribed genes (Fig. 2E). Although HU homologs have been reported to prefer to bind to AT-rich regions, we did not observe an obvious correlation between HU-binding sites and AT-rich regions in the pneumococcal genome (Supplementary Fig. 2). Taken together, ChIP-Seq analysis showed that HU binds with low intensity the whole chromosome, with a bias towards the origin of replication and at highly transcribed loci.

### Depletion of HU severely impacts growth and cell morphology

To be able to tightly control HU expression and investigate its role in chromosome dynamics, we replaced the native HU promoter with an anydrotetracyline (aTc) inducible promoter. Growing the cells in the absence of aTc led to a minor growth rate defect compared to the wild-type and cells grown in presence of the inducer (+ aTc) (Supplementary Fig. 3A). Phase-contrast and fluorescence microscopy of cells stained with DAPI revealed defects in cell morphology and the appearance of a small percentage of a-nucleated cells, although still compatible with growth (Supplementary Fig. 3B). We hypothesise that this approach was not enough to drastically reduce HU cellular levels, as also observed for *B. subtilis* (Karaboja and Wang 2022). Therefore, we fused Ptet-HU to superfolder GFP (Ptet-HU-sfGFP) to obtain a more unstable HU variant. In this case, removal of the inducer led to a drastic reduction of growth rate (Fig. 3A). Even in the presence of the inducer, the depletion strain showed a growth defect compared to the wild-type D39V. To confirm that the growth defect observed in absence of the inducer was solely due to reduced HU levels (p-HU, for pneumococcal HU), we complemented HU with a second copy of *E. coli α-*HU (e-HU), expressed ectopically under an IPTG inducible promoter. e-HU was able to restore growth, as well as restore the loss of nuceloids (Supplementary Fig. 3C-D).

**Figure 3.**
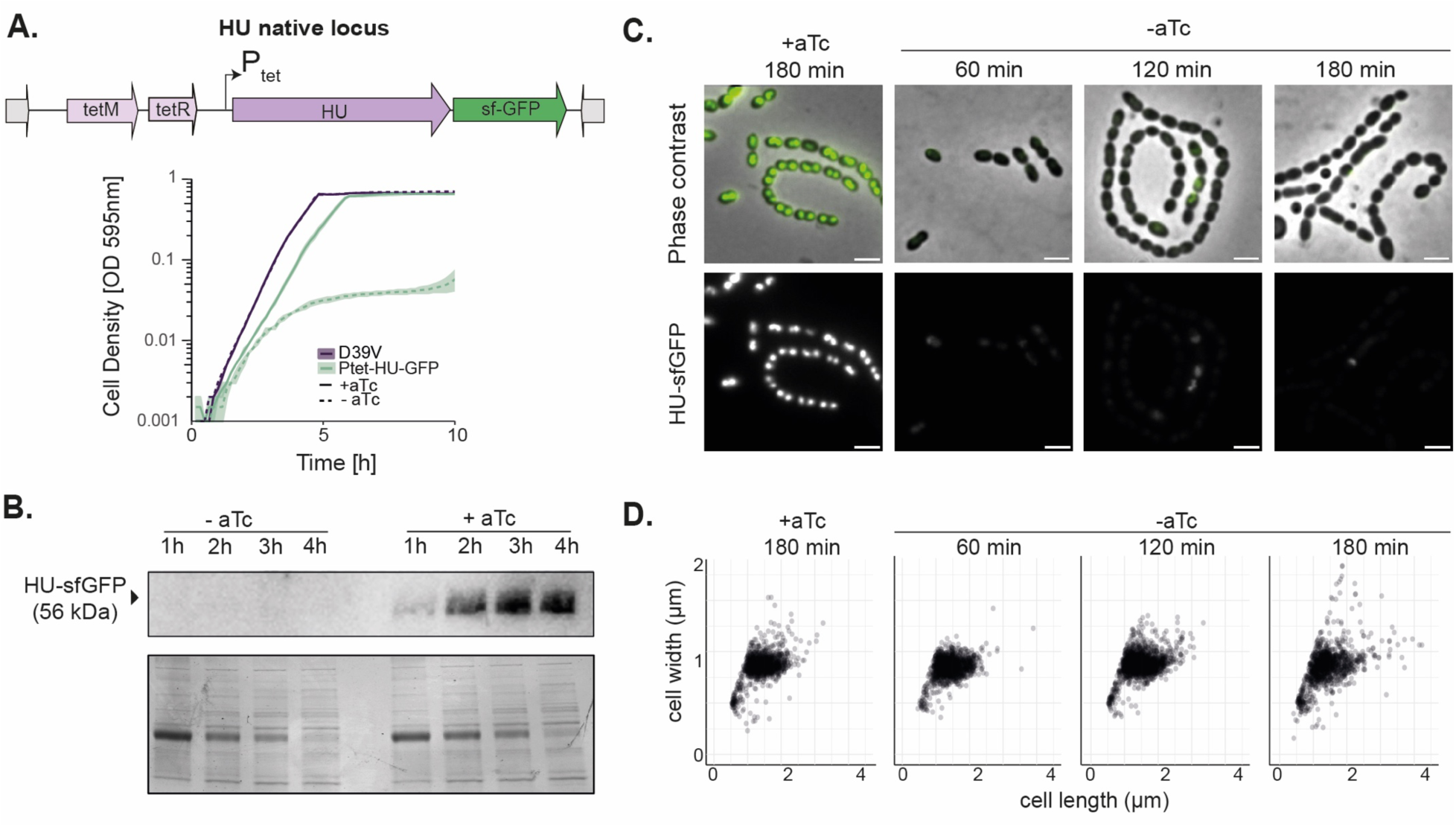
HU depletion kinetics and effects on cell morphology. **A.** Overview of HU depletion strategy. The native HU promoter P*hu* was replaced with a P*tet* promoter at the native locus. A selection marker (*tetM*) was added upstream, as well as *tetR* for aTc induction. Growth curves of cells with HU depletion (Ptet-HU-sfGFP) (-aTc) indicate a strong growth defect. **B.** Depletion kinetics of Ptet-HU-sfGFP by immunoblot (anti-GFP antibody) at one, two, three and four hours of depletion. Coomassie blue staining was used as a loading control (bottom). **C.** Depletion kinetics of Ptet-HU-sfGFP by fluorescence microscopy. HU-sfGFP expression is drastically reduced after one hour of depletion. Top: composite of HU-sfGFP and phase-contrast. Bottom: HU-sfGFP. Scale bar: 2 μm. **D.** Distribution of cell width and length of HU-depleted cells.

Western blot analysis using anti-GFP antibodies showed that depletion of HU was very efficient with significantly reduced HU-sfGFP after one hour and with a complete loss of signal after three hours in absence of inducer (Fig. 3B). Using fluorescence microscopy, we observed strains grown with and without aTc at three distinct time intervals to assess depletion kinetics and cell shape (Fig. 3C-D). Ptet-HU-sfGFP cells grown with aTc gave a bright signal that well distinguishes the pneumococcal nucleoid, which is lost after one hour of depletion, confirming the immunoblot observations (Fig. 3B). While there was no significant difference in the total cell size distributions (Kolmorogov-Smirnov test, p<0.05 for all combinations), a typical, severe morphological defect was visible; hallmarked by an increased irregularity in cell size within cell chains (Fig. 3C), and an increase in the percentage of extremely long cells (Fig. 3D, cell length > (mean+2*sd)_+aTc_, +aTc = 2.9%, -aTc: 60 min = 13 %, 120 min = 8.7 %, 180 min = 12%). These observations defined HU as an essential NAP, required for pneumococcal growth and cell shape in line with previous observations (Ferrándiz et al. 2018).

### Reducing HU levels alters nucleoid scaling dynamics

We next investigated nucleoid morphology upon HU depletion using wide-field fluorescence microscopy of cells stained with DAPI (Fig. 4A). Over time, in absence of aTc, the percentage of anucleate cells increases, being 10% at 1h (125 out of 1463 cells), 12.5% at 2h (250 out of 2268 cells) and 55% at 3h (851 out of 1547 cells) of depletion, suggesting a chromosome replication or segregation defect (Fig. 4B). The disruption of the chromosome integrity observed in a HU depletion strain was phenocopied by treatment with antibiotics affecting the cellular supercoiling state and DNA replication, as well as a *smc* deletion (van Raaphorst et al., 2017 and Fig. 4E-F). Indeed, cells depleted of HU were sensitised to treatment with sub-inhibitory doses (sub-MIC) of antibiotics that inhibit DNA gyrase and topoisomerase IV (Ciprofloxacin, CIP), and DNA replication (6-p-hydroxyphenylazo-uracil, HPUra) (Brown 1970). However, treatment with low doses of spectinomycin, which inhibits protein synthesis, did not show a synergistic effect with HU depletion (Supplementary Fig. 4).

**Figure 4.**
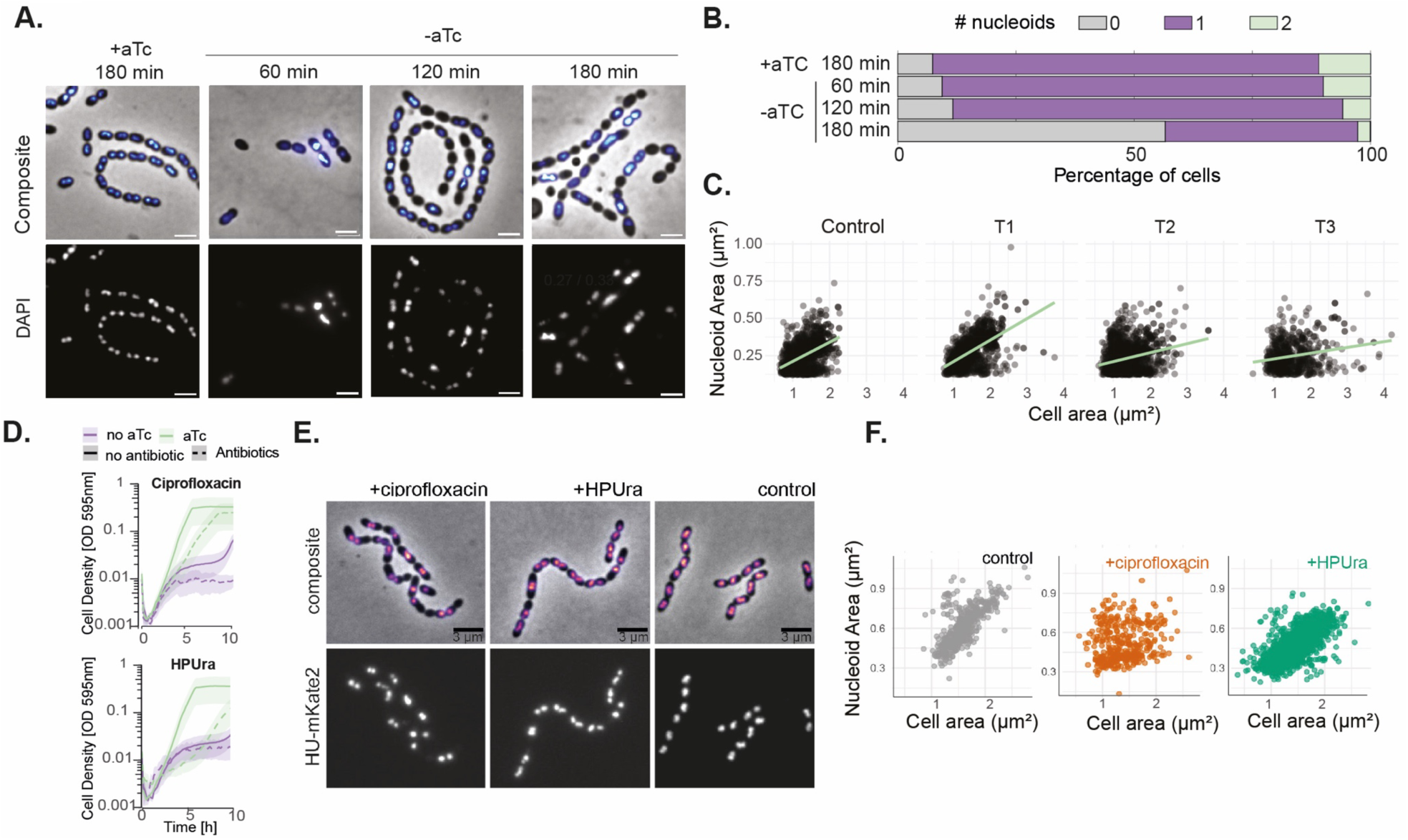
Depletion of HU strongly affects chromosome maintenance and NC scaling. **A.** Fluorescence microscopy of HU depleted cells stained with DAPI and imaged immediately afterwards. Scale bar: 2 μm. Cell area and nucleoid area were segmented from phase-contrast (cells) and fluorescence (nucleoids) with Oufti. **B.** Percentage of cells lacking a nucleoid increase with time (55% after three hours of depletion). **C.** Nucleoid to cell area (NC) for HU-depletion strain grown in presence (Control) and absence of aTc for one (T1), two (T2) and three hours (T3). **D.** Growth of HU-depletion strain in presence and absence of the inducer and with sub-MIC concentrations of HPUra (0.1 ug/mL) and ciprofloxacin (0.2 ug/mL) **E.** Representative images of cells carrying *hu::hu-mKate2* grown with sub-MIC concentrations of ciprofloxacin or HPUra, Scale bar: 3 μm**. F.** NC ratios are distorted in cells treated with ciprofloxacin but not in cells treated with HPUra.

In replicating wild-type pneumococci, the nucleoid is remarkably compact and robustly scales with cell size during growth, with a median nucleoid size of 0.5 µm^2^ and a nucleoid to cell ratio (NC) of 0.4. (Fig. 4C, first panel and F, control). After three hours of HU depletion, the NC ratio of cells that still have a nucleoid is drastically perturbed, with the appearance of larger cells with small nucleoids (Fig. 4C, Pearson’s R / median NC ratio: +aTc = 0.49 / 0.39, -aTc; 60 min = 0.57 / 0.37, 120 min = 0.27 / 0.33, 180 min = 0.21 / 0.32). To investigate what causes this NC perturbation, we measured the nucleoid size using HU-mKate2 as a nucleoid marker in cells treated with HPUra and CIP. Blocking DNA replication using sub-MIC concentrations of HPUra led to anucleate cells, however, the nucleoid scaling ratio stayed intact, confirming that NC scaling is independent of DNA replication (Gray et al. 2019) (Fig. 4D-F). Cells lacking *smc* showed a phenotype very similar to HPUra treatment (van Raaphorst et al., 2017). Interestingly, when cells were grown in presence of sub-MIC concentrations of CIP, the relationship of the nucleoid to cell size is lost, due to aberrant cell and nucleoid sizes (Fig. 4D-F). Overall, we showed that depletion of HU severely impacts chromosome maintenance and disrupts the robust nucleoid scaling with cell size. Our data suggests that these phenotypes may be attributed to significant defects in chromosome transitions, including DNA supercoiling and DNA repair, resulting from HU depletion.

### HU promotes the higher organisation of the pneumococcal chromosome

To investigate the exact three-dimensional organisation of the pneumococcal nucleoid and the impact of HU in chromosome conformation, we performed Hi-C in HU-depletion strain. Cells were grown with and without aTc for three hours until exponential phase was reached (Fig. 5A, B). Samples were processed using Hi-C, and DNA was sequenced and analysed to produce contact maps. In cells grown in presence of aTc and expressing HU (Fig. 5A), the contact map revealed, besides the expected diagonal signal resulting from contacts between neighbouring loci, faint secondary diagonal. This secondary diagonal is a common feature of bacterial genomic Hi-C maps and reflects enrichments in contacts between chromosomal arms and mediated by condensins (X. Wang et al. 2015; Marbouty et al. 2015; Le, Laub, 2013, Marbouty et al. 2017; Umbarger et al. 2011). The presence of other structural signatures such as self-interacting or large domains are difficult to assess in the present dataset, either because of a lack of resolution, or because the exponentially growing cells lack such structures.

**Figure 5.**
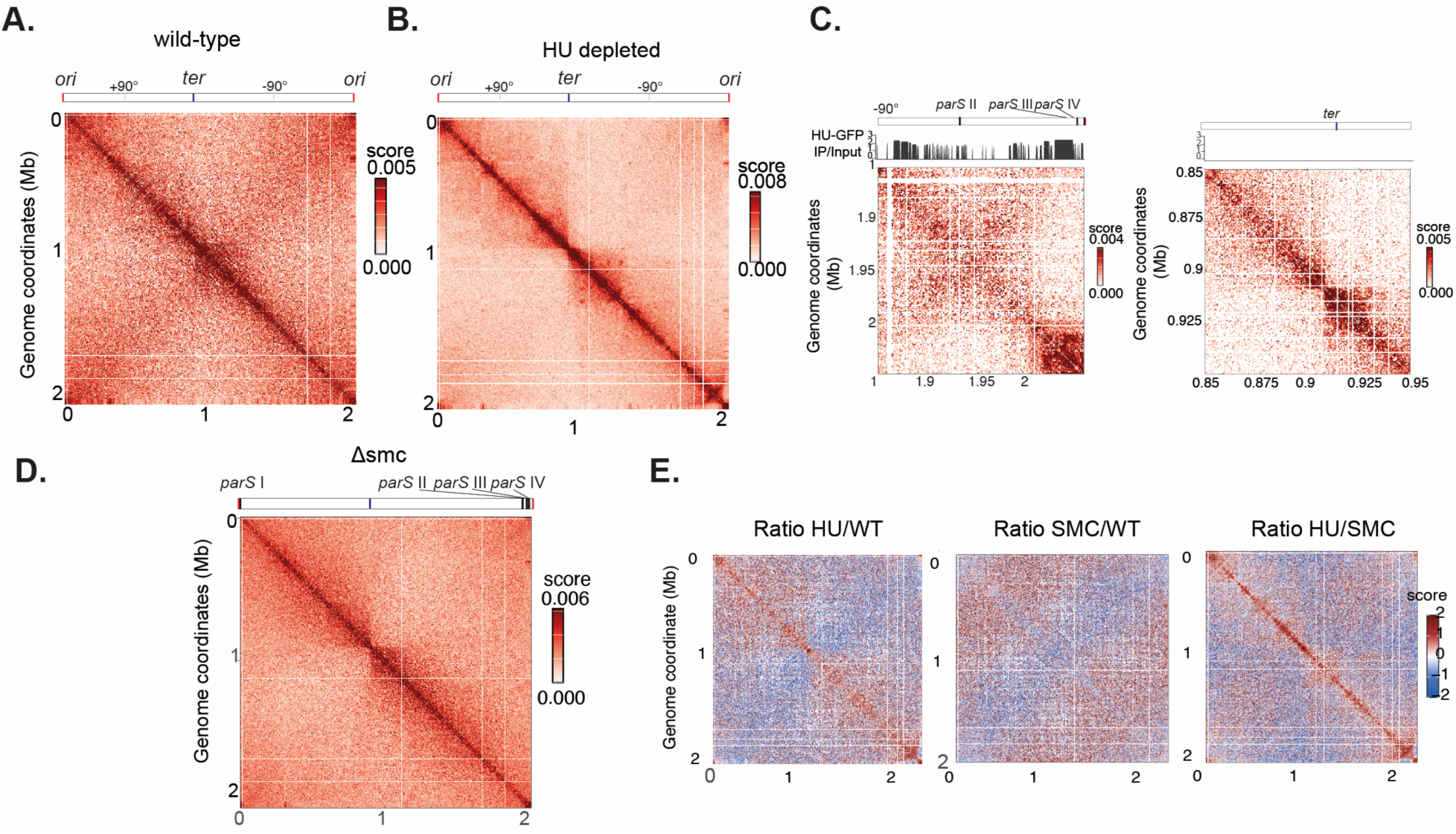
HU and SMC promote long-range chromosome folding. **A.** Normalised Hi-C contact maps of wild-type D39V cells at 5 kb resolution Position of the origin and terminus of replication are depicted. **B.** Normalised Hi-C contact maps of HU-depletion strain grown in absence of aTc (three hours depletion) at 5 kb resolution. **C.** Magnification of the origin on the left replichore (upper panel) and terminus regions (100 kb). *Ori, ter* and *parS* binding sites are depicted. **D.** Normalised map of Δ*smc* at 5 kb resolution. **E.** Differential contact maps corresponding to the log2 ratio of Hi-C interactions between WT and cells lacking HU or SMC, and between cells lacking HU and SMC. The blue/red colour scale reflects the enrichment in contacts in one population with respect to the other.

In absence of HU, the normalised contact maps showed significant differences compared to WT conditions. First, contact matrices showed a reduction in long-range contacts (1 Mb) compared to the wild-type strain, concomitant with an increase in short-range contacts along the chromosome (Fig. 5A and B). Secondly, replichores trans contact were strongly attenuated, with the origin of replication clearly partitioning the chromosome. Interestingly, local structuration of the chromosome also emerged in absence of HU, including dots in the maps likely suggesting the presence of DNA loops, as well as DNA stripe or ter-proximal “hairpin” structure on the left replichore, of ∼100 kb (Fig. 5B and 5C). Magnification of 100 kb surrounding the *ori* on the left replichore (Fig. 5C), underlined a dense HU binding in this region, overlapping the *parS*-SMC loading sites II in correspondence of the hairpin and III and IV closer to the *oriC* (Minnen et al. 2011). At *ter* (100kb), the conformational signature reflects the increased cis-contacts between the two replichores due to loss of cohesion (Fig. 5C). To compare the impact of HU depletion with the absence of the SMC condensing known in other species to be responsible for replichore bridging, we performed Hi-C in a Δ*smc* mutant (Fig. 5D). As expected, the Hi-C contact map confirmed a role of SMC in bridging the pneumococcal chromosome replichores. Remarkably, reduction of long-range contacts obtained in absence of SMC is similar to what was observed in the absence of HU (Fig. 5D). The ratio of normalised contact maps between wild-type, HU-depleted and Δ*smc* cells nevertheless pointed at differences between HU and SMC depleted chromosomes (Fig. 5E). Indeed, whereas HU depletion led to changes in the local folding of chromosomes, the loss of SMC had little impact on that organization. In particular, the segmentation at the level of the *ori* and overall, the enrichment in short-range contact was less pronounced in absence of *smc*, and the ter-proximal hairpin was absent. Altogether, these findings support a major role of HU in promoting long-range contacts and maintaining the conformation of the origin domain.

### HU is required for replication initiation and replisome localisation

To investigate whether the lack of HU directly impaired DNA replication, we calculated an index of replication (iRep) based on the sequencing coverage trend of exponentially growing cells and used it to infer replication rates. Cells were grown up to three hours without aTc (HU-depleted cells) and three hours with the inducer. Genomic DNA was extracted, processed, sequenced, and analysed using iRep (Brown et al. 2016). For cells grown in presence of aTc, reads mapped to the wild-type genome (D39V) and plotted over the correspondent chromosomal position showed a typical profile of exponential growth, with a iRep value (normalised ratio of *ori-*to*-ter* coverage) of 2.4 (Fig. 6A). This value is very similar to that of D39V wild-type strain (Slager et al. 2014), indicating that the HU-depletion strain grown in presence of the inducer replicates at a similar rate. Upon HU depletion, the profile became flat, and the iRep value dropped to 1.77 at one hour, 1.28 at two hours and 1.17 at three hours, indicating that the HU-depleted cells progressively stopped replication.

**Figure 6.**
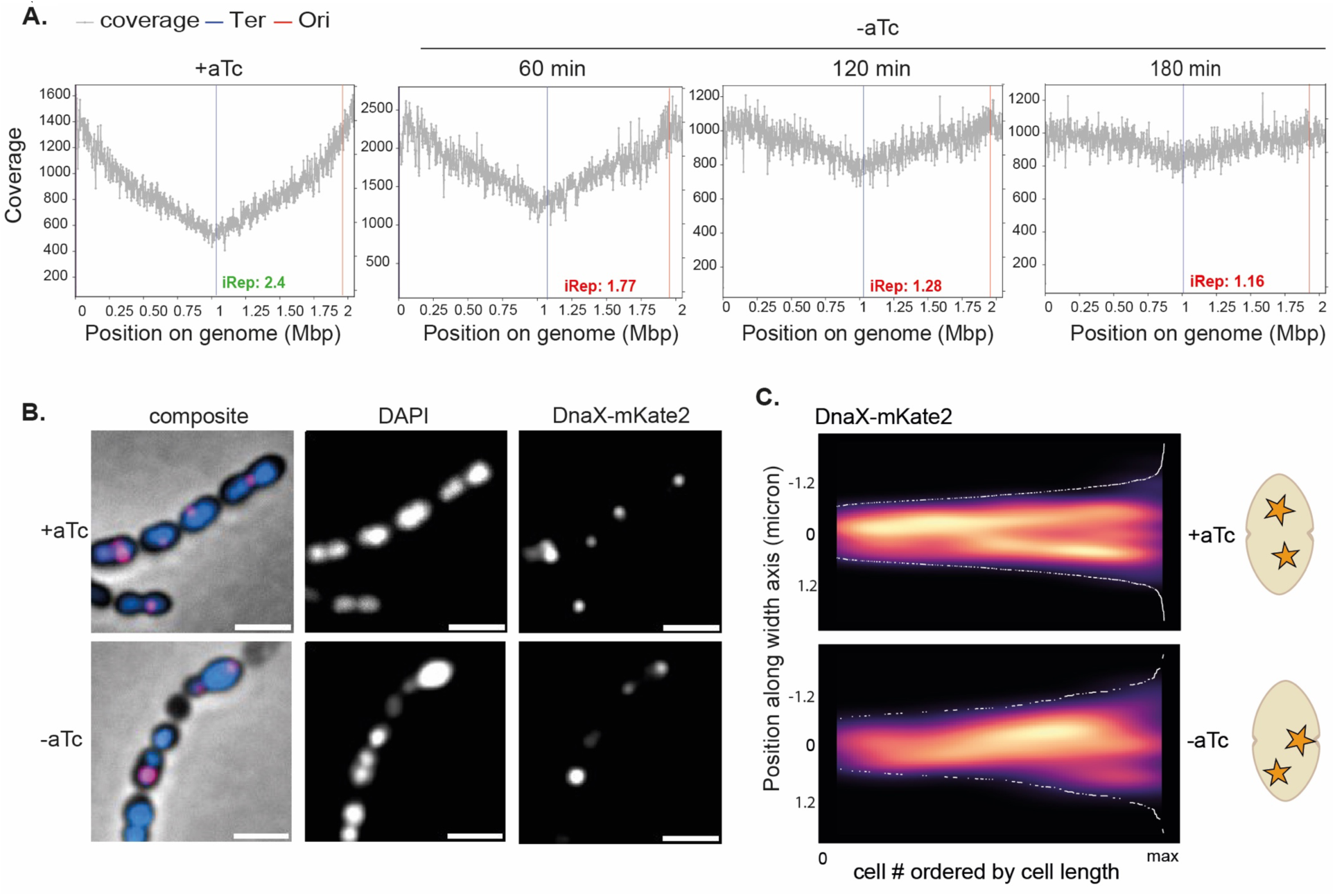
HU-depleted cells progressively stop replication. **A.** Genome-wide marker frequency analysis. Sequencing reads coverage over the genome sequence of exponentially growing cells in presence of aTc (on the left) and at one, two, and three hours of depletion. The replication rate of the population is indicated by the ratio of coverage at the origin ("peak") to the terminus ("trough") of replication. The origin and terminus position are determined based on cumulative GC skew. **B.** DnaX-mKate2 localization in cells depleted for HU (-aTc) for 180 minutes and non-depleted cells (+aTc). Left to right: composite (phase: grey, DAPI: blue, DnaX-mKate2: pink), DAPI, DnaX-mKate2. **C.** Kymograph (left) with graphical representation (right). In cells with HU, DnaX is localized at mid-cell and migrates towards the new septa during growth (+aTc, total cells = 313). In cells depleted for HU for 180 minutes (-aTc, total cells = 705), DnaX is mislocalized.

To follow the DNA replication machine (replisome) in live cells, we fused the clamp loader DnaX with a red fluorescent protein (strain *dnaX::dnaX-mKate2*) and imaged the dynamics of the replisome by fluorescence microscopy (Fig. 6B). Fluorescence microscopy showed enriched signals as bright diffraction -limited spots in HU-depletion strain grown in presence and absence of the inducer, indicative of active replication. Interestingly, HU depletion decreased the percentage of actively, normally replicating cells (defined as cells with 1-2 DnaX foci) from 66% percent to 49% (Supplementary Fig. 5A). In wild-type cells, replication forks localise at mid-cell and migrates towards the new septa during growth, while in absence of HU are often mislocalized (Fig. 6C), reminiscent of cells with impaired chromosome integrity (van Raaphorst, Kjos, and Veening 2017). Together, these results show that, in the absence of HU, cells fail to initiate replication rounds. When the DNA replication machinery is assembled, further initiation, localisation and progression of the replication fork is impaired.

### HU genetic interactions and a model for chromosome organisation

To gain further insights into the cellular processes to which HU contributes, we exploited the recently generated dual-CRISPRi-seq dataset PneumoGin (Dénéréaz et al. bioRxiv). By simultaneously downregulating the expression of two genes or operons, a dual-sgRNA CRISPRi library was used to screen for genetic interactions in *S. pneumoniae*. These interactions can be negative (simultaneous repression of two genes worsen the growth defect) or positive (simultaneous repression restore growth). By setting a threshold of epsilon of 2, 10 genes or operons are found to be synthetic lethal with HU (Fig. 7A, data available on pneumoGIN). Most of the genes are involved in replication and repair (COG:L) and cell cycle control (COG:D) pathways. These include the DNA topoisomerase I (*topA),* the helicases *pcrA* and the resolvase *recU.* Proteins required for positioning of the replication machinery and chromosome segregation such *scpA*/*B,* the integral components of the Smc/ScpAB complex, are also synthetic lethal with HU. In addition, knock-down of genes involved in cell size determination (*pgm*) and cell wall biosynthesis (*pbp1a*) worsen a HU phenotype, in line with the deleterious morphology perturbations observed upon HU depletion. Genes with unknown functions (SPV_0750 and SPV_0761) are also part of the HU interaction network, highlighting that HU essentiality might be linked to yet unexplored cellular functions. No positive interactions were identified, indicating that the defects of reduced HU levels cannot be restored by the knock-down of any pneumococcal gene.

**Figure 7.**
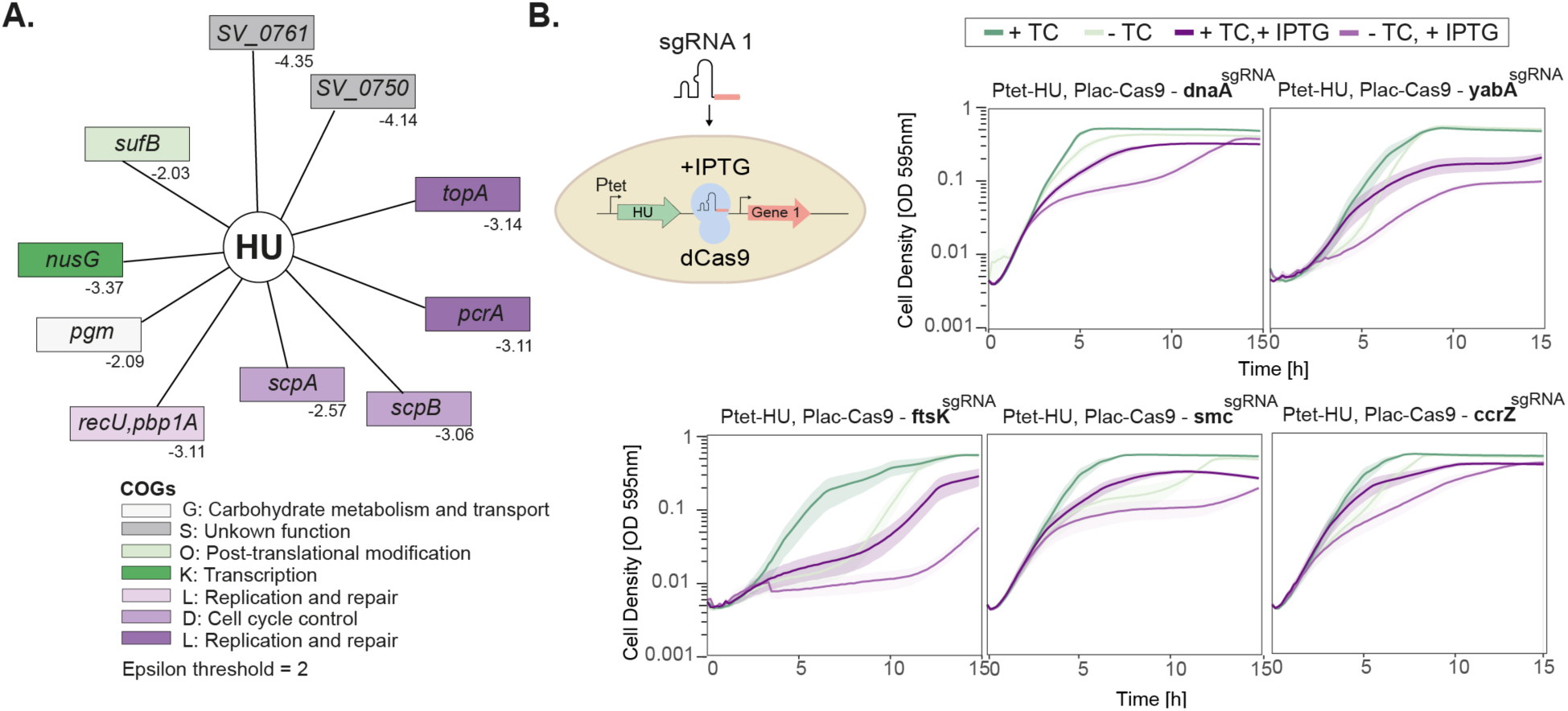
HU is a major structural component essential for chromosome conformation, DNA replication, and chromosome segregation. **A.** Genetic interaction network of HU as determined by dual-CRISPRi-Seq (Dénéréaz et al. bioRxiv). Epsilon values are indicated, and a threshold of 2 is set and defined as the added fitness effect of the knockout of two genes on top of the fitness effects caused by the knockout of those same two genes individually. **B.** Growth of knock-down strains obtained by CRISPRi for the DnaA, YabA, CcrZ and SMC genes in a HU-depletion background. Absence of aTc leads to HU depletion, while presence of IPTG leads to expression of Cas9 and targeted knock-down of each gene.

To investigate the synthetic lethality of genes involved in DNA replication (DnaA, YabA and CcrZ) and chromosome segregation (SMC, FtsK), we constructed CRISPRi knock-downs and measured their growth (Fig. 7B). To do so, we inserted an IPTG-inducible dCas9 and single guide RNA (sgRNA) targeting each gene of interest in the Ptet-HU depletion strain. Notably, when growing the cells in absence of HU (- aTc) and inducing *cas9* expression (+ IPTG), we confirmed the synthetic lethality for all genes tested. These interactions enabled us to outline a model of pneumococcal chromosome organisation and segregation, incorporating all yet identified partners. In wild-type cells, the nucleoid alternates dynamically between “pull” and “twist” morphologies. The initiator (DnaA) and inhibitor (YabA) of DNA replication, the former controlled by CcrZ, determine the timing of replication and replisome progression. ParB binds to ParS sites and recruits the SMC complex, which together with FtsK and RocS, segregate the newly replicated chromosomes. In absence of HU, the supercoiling state of the cell is perturbed, as well as the high-order chromosome folding and the origin of replication. As a result, cells will either have a condensed nucleoid in a larger cell or lose it, either way leading to cell death. In this model, HU plays a central role in maintaining the supercoiled state and building the conformation required for initiation and progression of DNA replication and chromosome segregation.

## Discussion

Most diderm bacteria carry multiple nucleoid associated proteins (NAPs) which are functionally redundant (Dillon and Dorman 2010). In monoderm bacteria, this is not the case, and *S. pneumoniae* is an extreme example. Although new putative classes of NAPs have not been investigated in pneumococcus, there are no homologues of the IHF, H-NS, or FIS families, and HU is the only confirmed NAP. This lack of redundancy positions HU as a single, essential protein and makes the pneumococcus a particularly interesting model organism to study its role. Furthermore, due to its high expression (Aprianto et al. 2018) and overlap with the nucleoid, HU serves as an ideal marker for tracking the dynamics of the pneumococcal nucleoid. In this study, PALM experiments revealed an extremely compact nucleoid, with HU constantly moving along the chromosome (Fig. 1).

Drastically reduced levels of HU resulted in pleiotropic phenotypes, severely impairing cell shape and nucleoid maintenance. After three hours of depletion, more than one in two cells lost the nucleoid. This strong segregation defect partly answers one of the long-standing questions about how the pneumococcus manages to segregate its chromosome while most of the known factors (SMC, ParB, FtsK) can be eliminated with only mild phenotypes (van Raaphorst, Kjos, and Veening 2017; Kjos and Veening 2014) and places it as an essential factor for successful segregation.

Pneumococcal HU is required to maintain the supercoiling state and counteracts the effects of a less active gyrase (Ferrándiz et al. 2018). We showed that inhibiting gyrase and topoisomerase IV with a fluoroquinolone resulted in a nucleoid-to-cell ratio that mimicked HU depletion, supporting a critical role of HU in supercoiling, a primary reason for its essentiality. Although cells lacking HU do not replicate, HU is not required for the assembly of the replication complex. Given the denser binding of HU in proximity of the replication origin and the structural perturbations observed in a HU-depletion strain, we suggest that both supercoiling and architectural defects are responsible for the replication failure. This is also supported by the mislocalisation of the replication fork that we measured in absence of HU.

Hi-C data showed that HU-depleted cells also have lost the interarm chromosome interaction that is characteristic of SMC-mediated chromosome segregation (X. Wang et al. 2015; Marbouty et al. 2015; X. Wang et al. 2017; W. Wang et al. 2011; Tran, Laub, and Le 2017). We hypothesise that early in the cell cycle, HU is recruited at the *oriC* proximal region to build the conformation required for DNA replication and SMC-mediated chromosome segregation. A more precise view of the timing of events could be obtained by repeating these measurements at shorter time points after removal of the inducer, or to synchronise cells and perform single-cell measurements. We however speculate that the arrest of DNA replication and DNA segregation occur concomitantly, since SMC is not known to require active replication to process DNA. While the impact of HU deletion on chromosome conformation varies among bacterial species (Ponndara et al. 2024), the decrease in long-range contacts that we observed has also been noted in *E. coli* and is similar to what is observed in the absence of MukBEF (the *E. coli* condensin), suggesting a cooperation between condensin complexes and HU to promote the higher-order organisation of the chromosome (Lioy et al. 2018). This interplay has also been identified in *Streptomyces venezuelae (Szafran et al. 2021)*, and we suggest that it could also be the case for *B. subtilis*. In support of this, we find a strong negative genetic interaction between HU and the SMC complex by dual CRISPRi-seq (Fig. 6). We speculate that the hairpin observed in proximity of the *ori*, encompassing one *parS* site, could be explicative of the early loading of ParB/SMC (Brandão et al. 2021, X. Wang et al. 2017, Brandão et al. 2021), and a failure in proceeding through the chromosomal arms leading to a successful segregation.

Whether the role of HU in DNA replication and segregation involves direct interaction and modulation of DnaA and/or SMC/ParB activity is yet to be elucidated. However, the complementation obtained with *E. coli* HU (59% sequence homology) suggests that unlikely all interactions with homologous proteins are correctly maintained. This rather suggests a structural role required to support replication and the initial steps of loop formation by ParB/SMC. However, to investigate this further, biochemical, and structural studies are needed to examine HU interaction partners and binding at *ori*.

Dual-CRISPRi-Seq identified the genetic partners of HU at the genome-wide level (Dénéréaz et al. bioRxiv). While we cannot exclude the possibility that, given the partial down-expression of HU, other important genetic interactions may be missing, this work has provided a comprehensive view of the cellular pathways to which HU is integrated. Although multiple replication and partitioning factors are responsible for cell cycle progression, HU is the only protein with a structural role yet known in pneumococcus, strictly required for higher-order chromosome conformation and thus DNA replication and chromosome segregation.

## Data availability

Sequencing data from ChIP-seq, Hi-C experiment and Illumina whole genome sequencing are available in the Sequence Read Archive under BioProject accession code SUB14752130. ChIP-Seq data obtained for HU-GFP and RpoB-GFP, as well as Hi-C contact matrices, are available for visualisation in Pneumo browse 2.0 (doi: https://doi.org/10.1101/2024.08.07.606308). R scripts used for image analysis are made available on GitHub: https://github.com/veeninglab/HUproject.

## Acknowledgements

We thank Stephan Gruber for useful discussions. Thanks to Vincent de Bakker for help with the analysis of the RpoB ChIP-Seq data. We thank the Genome Technologies Facility (GTF) department in Lausanne for the sequencing and library preparation of the ChIP-Seq experiment and the sequencing platform facility of Institut Pasteur for Hi-C sequencing.

## Authors Contributions

M.V.M and R.V.R wrote the paper with input from all authors. M.V.M, R.V.R, L.M, F.B, B.W, and J.S performed the experiments. M.V.M, R.V.R and J.W.V designed, analysed, and interpreted the data. All authors reviewed and approved the final version of the manuscript.

