## Supplementary figures and images for "HU promotes higher-order chromosome organisation and influences DNA replication rates in *Streptococcus pneumoniae*"

### Supplementary Figure 1

**Supplementary figure S1.**

**A.**

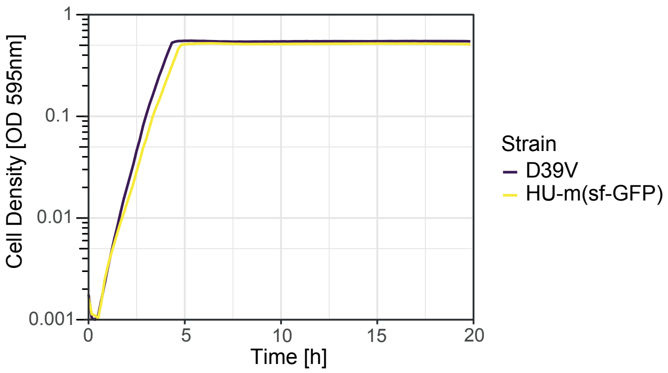


**B.**

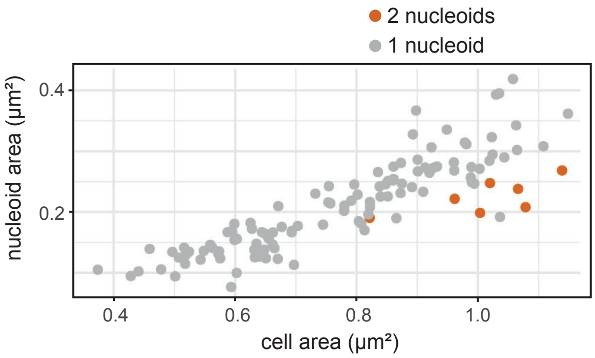


**C.**

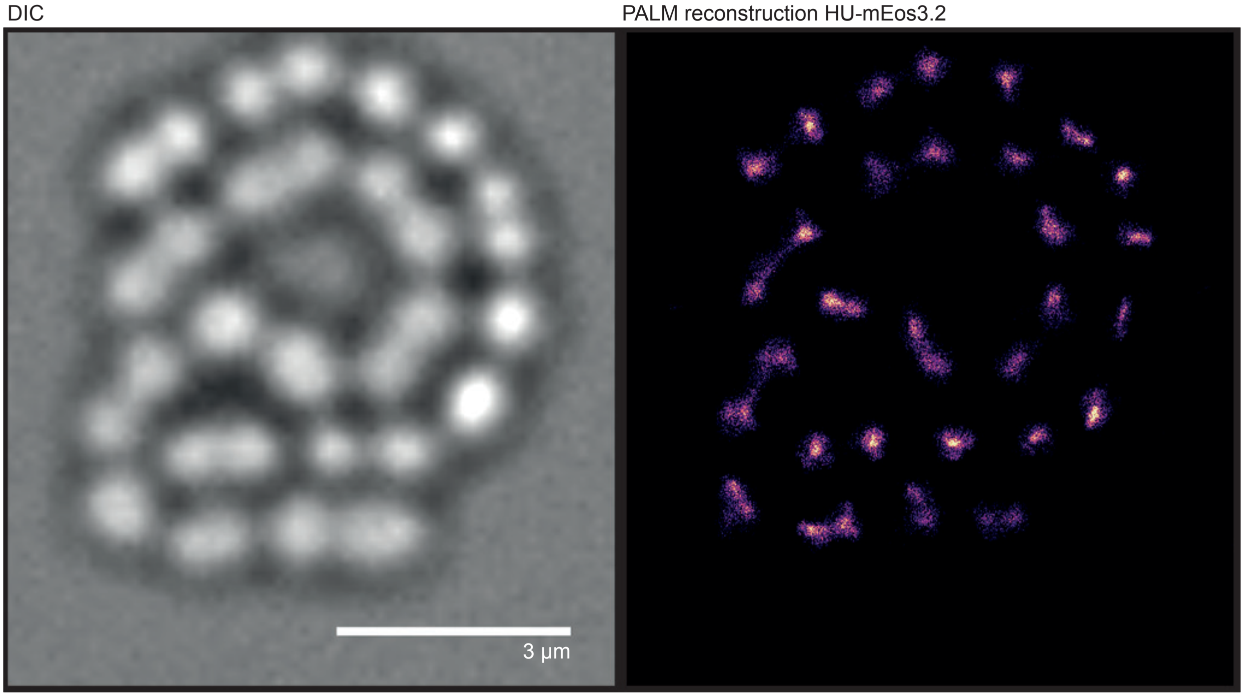

### Supplementary Figure 2

#### Supplementary figure S2.

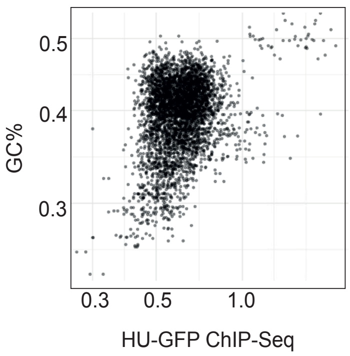

### Supplementary Figure 3

**Supplementary Figure 2.**

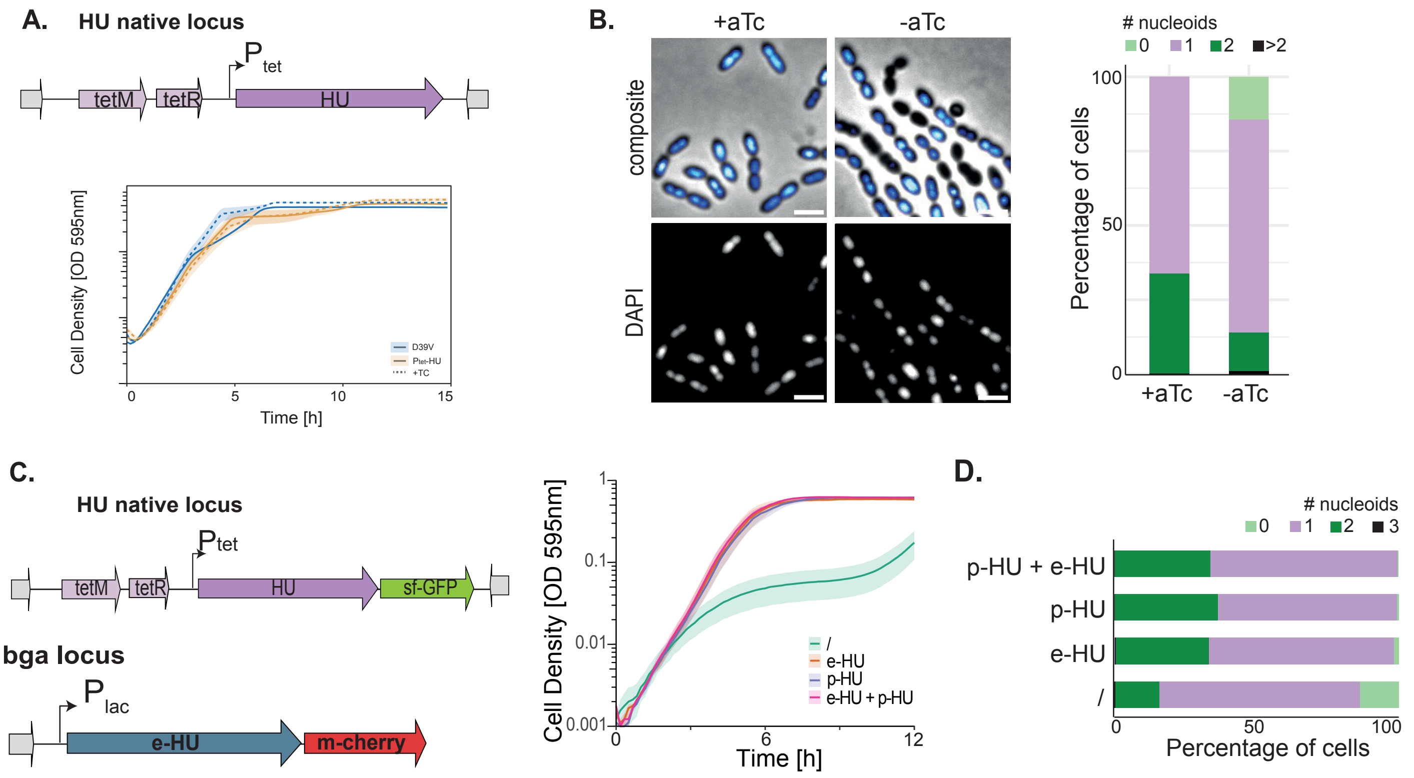

### Supplementary Figure 4

**Supplementary figure S4.**

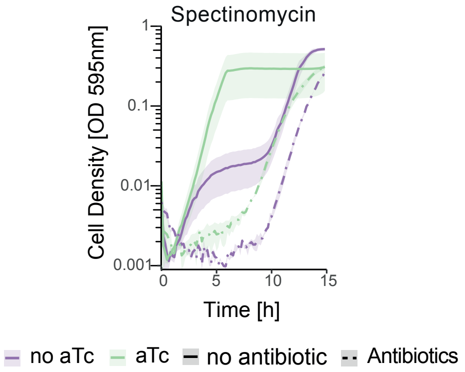

### Supplementary Figure 5

#### Supplementary figure S5.

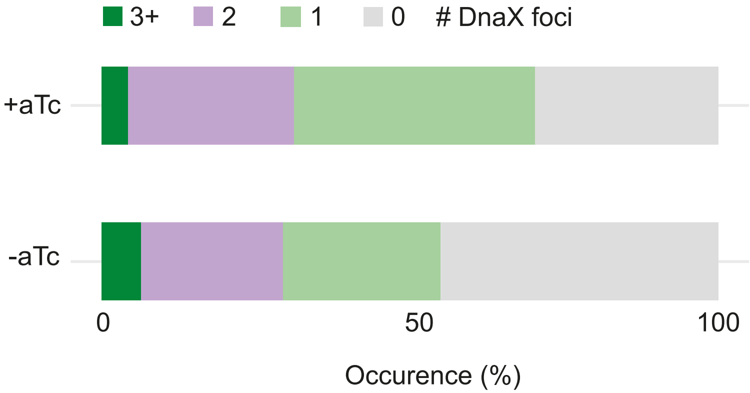
